# Analyzing globin gene variation in gnomAD: implications for variant interpretation in hemoglobinopathies

**DOI:** 10.64898/2026.09.14.750622

**Authors:** Kontopoulou Mikaella, Xenophontos Maria, Lederer Carsten Werner, Stephanou Coralea, Kountouris Petros

**Author notes:** Joint last authorship (Stephanou Coralea, Kountouris Petros). Corresponding author Kountouris Petros. **Authors’ email addresses**: Kontopoulou Mikaella, Xenophontos Maria, Lederer Carsten Werner, Stephanou Coralea, Kountouris Petros.

## Abstract

**Background:** Given the augmented sampling and ancestry stratification of the Genome Aggregation Database (gnomAD) v4.1 compared to v2.1.1, population evidence for globin gene variants was assessed for changes in the American College of Medical Genetics and Genomics and the Association for Molecular Pathology (ACMG/AMP) criteria fulfillment, which can alter downstream variant interpretation.

**Methods:** Variants and constraint metrics for α-globin locus genes *HBA1*, *HBA2*, and *HBZ*, and β-globin locus genes *HBB*, *HBD*, *HBG1*, *HBG2*, and *HBE1*, were extracted from gnomAD v2.1.1 and v4.1 callsets, harmonized, matched to ClinVar annotations, and evaluated against BA1, BS1, BS2, BS2_Supporting, and PM2_Supporting ACMG/AMP criteria using Clinical Genome Resource Hemoglobinopathy Variant Curation Expert Panel thresholds.

**Results:** Globin gene variants increased fourfold across gnomAD releases, with *HBB* contributing the largest share. In v4.1, 68.7% of quality-passing variants were ancestry-specific, 14.4% overlapped ClinVar and PM2_Supporting fulfillment rose from 81.5% to 93.5%. Among variants present in both releases, ACMG/AMP population-criterion assessments were largely concordant, with conflicting combinations in <1%.

**Conclusions:** Leveraging augmented sampling and improved ancestry representation, gnomAD v4.1 increased globin gene variant discovery, particularly for rare variants, enabling more precise allele frequency estimates and robust assessment of BA1, BS1, and PM2_Supporting, although criterion fulfillment remains release– and callset-specific.

## Introduction

Hemoglobinopathies comprise a heterogeneous group of inherited disorders resulting from pathogenic variants in coding and regulatory elements of the α-(NG_000006) and β-globin (NG_000007) gene clusters [1]. Collectively, they are the most prevalent group of monogenic disorders worldwide, with a distribution shaped by selection in malaria-endemic regions and contemporary population movement that redistributes affected individuals across geographic and socioeconomic divides [2,3]. In regions with historically high malaria transmission, pathogenic globin gene variants reach atypically high frequencies, despite detrimental consequences of homozygous or compound heterozygous genotypes. Their persistence at elevated frequencies reflects balanced polymorphism, whereby pathogenic globin gene variants are maintained by a heterozygous selective advantage against severe malaria [4]. Based on the founder effect and a high level of allelic heterogeneity, different affected regions and populations may thus be characterized by an elevated frequency of distinct sets of pathogenic globin gene variants [5].

Diagnostic sequencing reveals an expanding catalog of distinct globin gene variants and complexity beyond the globin loci [3]. Clinical significance depends not only on the intrinsic effect of an individual variant but also on allelic configuration, co-inherited globin gene variants, and a growing list of genetic disease modifiers [5,6]. This complicates genotype-phenotype inference and renders variant interpretation central to accurate molecular diagnosis, carrier screening, reproductive counseling, and genotype-informed management [6,7].

Standardized interpretation is therefore essential. The joint guidelines of the American College of Medical Genetics and Genomics and the Association for Molecular Pathology (ACMG/AMP) provide a structured framework for evaluating and classifying sequence variants as pathogenic, likely pathogenic, of uncertain significance, likely benign, or benign [7]. These guidelines specify how diverse evidence types, including population data, segregation, functional assays, case observations, computational predictions, and other lines of evidence, should be weighted and integrated [7,8].

Within this framework, population allele frequencies constitute a core line of evidence. For Mendelian disorders, observed frequency must be compatible with disease prevalence, penetrance, and mode of inheritance [9]. Variants exceeding a maximum credible allele frequency [9] provide stand-alone or strong benign evidence (BA1 and BS1 ACMG/AMP criteria), while variants absent or extremely rare in reference populations may contribute supporting pathogenic evidence (PM2_Supporting ACMG/AMP criterion) [7]. Observations of homozygotes in reference population datasets (BS2 and BS2_Supporting ACMG/AMP criteria) further inform benign classification when disease penetrance is high [7].

Applying these criteria to hemoglobinopathies requires careful calibration, as use of overly simplistic frequency thresholds risks misclassification; either by classifying genuinely pathogenic, population-enriched variants as benign or inappropriately considering underrepresented benign ancestry-specific variants as pathogenic. To address this, the Clinical Genome Resource (ClinGen) Hemoglobinopathy Variant Curation Expert Panel (VCEP) has specified hemoglobinopathy-calibrated allele frequency thresholds and structured guidance for tailoring the ACMG/AMP criteria to globin gene variants in *HBA1*, *HBA2*, and *HBB* [5,10].

Population evidence for globin gene variants is anchored to the publicly available reference population dataset of the Genome Aggregation Database (gnomAD), which aggregates large-scale exome and genome sequencing data and reports ancestry-stratified allele frequencies and gene-level constraint metrics [11]. The latest release (v4.1) augments sample size, improves ancestry stratification, and implements joint exome-genome variant calling [11]. These improvements increase statistical power for rare variant detection and reduce uncertainty in population-specific frequency estimates, thereby enhancing the robustness of population criteria assessment. Current ClinGen guidance designates gnomAD v4.1 as the primary reference population dataset for the application of frequency criteria by the ClinGen Hemoglobinopathy VCEP and all other VCEPs [10]. Adoption of v4.1 is therefore embedded within the evolving interpretive framework applied to globin genes. Given the central role of allele frequencies in BA1, BS1, and PM2_Supporting application, augmented sampling may change whether these criteria are met, thereby influencing variant interpretation outcomes, particularly for variants that lie near the pathogenic-benign interpretive boundary in the absence of additional supporting data [12]. Given these potential consequences of the gnomAD version transition, a systematic assessment of its effect on globin gene variant interpretation based on population criteria is imperative.

Addressing this need, this study compares globin gene variation between gnomAD v2.1.1 and v4.1, quantifies changes in variant detection and allele frequency estimates, and evaluates the resulting impact on ACMG/AMP population criterion fulfillment using the ClinGen Hemoglobinopathy VCEP specifications. The objective is to evaluate how updated population data influence ACMG/AMP population criterion fulfillment for globin gene variants, with specific attention to variants shared across gnomAD releases.

## Materials and Methods

### Data acquisition

Constraint metrics and variants for *HBA1* (HGNC:4823), *HBA2* (HGNC:4824), *HBB* (HGNC:4827), *HBD* (HGNC:4829), *HBE1* (HGNC:4830), *HBG1* (HGNC:4831), *HBG2* (HGNC:4832), and *HBZ* (HGNC:4835) were extracted from the exome, genome, and joint callsets of gnomAD v4.1 and the GRCh38 liftover exome and genome callsets of gnomAD v2.1.1 (accessed 28 May 2025) [13]. v4.1 includes 730,947 exomes and 76,215 genomes; v2.1.1 includes 125,748 exomes and 15,708 genomes [13]. Variants were harmonized across releases and callsets by chromosome, position, reference allele, and alternate allele, then integrated with ClinVar annotations (accessed 10 October 2025), including pathogenicity classifications and associated review status [14].

### Variant characterization and comparative analyses

Gene-level constraint metrics were evaluated to contextualize gene-specific tolerance to variation and to assess whether increased sampling in gnomAD v4.1 altered inferred selective constraint relative to v2.1.1. Loss-of-function (LoF) observed/expected upper bound fraction (LOEUF), along with synonymous and missense constraint z-scores, were extracted from release-specific gnomAD summary tables [13,15]. LOEUF reflects tolerance to LoF variation; values below 0.6 indicate increased constraint and reduced tolerance [13]. Missense and synonymous constraint z-scores represent standardized deviations of observed from expected variant counts, with z≥3.09 indicating reduced tolerance to variation [13].

In parallel, variant presence and gnomAD quality assessment were compared across releases, callsets, and genes to quantify differences in variant detection and classification. Quality status was defined solely by gnomAD quality assessment annotations, without additional filtering. Quality assessment procedures in gnomAD are applied independently to callsets, with release-specific updates to variant-calling pipelines and filtering thresholds [13].

Unless otherwise specified, downstream analyses were restricted to quality-passing variants from the gnomAD v4.1 joint callset. A cross-reference with ClinVar variants was performed to estimate the proportion of globin gene variation not captured in a clinical variant resource. Quality-passing variants in v2.1.1 and v4.1 were evaluated for fulfillment of ACMG/AMP criteria BA1, BS1, BS2, BS2_Supporting, and PM2_Supporting using ClinGen Hemoglobinopathy VCEP thresholds (Table 1) [5,7]. Although Hemoglobinopathy VCEP specifications formally apply to *HBA1*, *HBA2*, and *HBB*, additional globin genes were analyzed to characterize locus-wide population frequency patterns and contextualize gene-specific criterion fulfillment. Variants meeting criterion combinations with conflicting interpretive implications were identified. For v2.1.1, criteria were evaluated using exome and genome data, prioritizing the callset with the higher population maximum filtering allele frequency (PopMax AF; defined as the highest filtering allele frequency observed among genetic ancestry groups, with 95% confidence interval) when present in both [16].

**Table 1.** ACMG/AMP variant interpretation criteria specified for globin genes^5,7^.

| **Criterion** | **Hemoglobinopathy VCEP Definition** | **Evidence Classification** |
| --- | --- | --- |
| **BA1** | Population maximum filtering allele frequency >0.5% in reference population dataset, with $\geq 5$ observed variant alleles and $\geq 2000$ alleles evaluated | Stand-alone evidence supporting a benign classification |
| **BS1** | Population maximum filtering allele frequency between 0.1% and 0.5% in reference population dataset, with $\geq 5$ observed variant alleles and $\geq 2000$ alleles evaluated | Strong evidence supporting a benign classification |
| **BS2** | Variant observed in the homozygous state in $\geq 2$ individuals without reported disease in the reference population dataset | Strong evidence supporting a benign classification |
| **BS2_Supporting** | Variant observed in the homozygous state in 1 individual without reported disease in the reference population dataset | Supporting evidence of benign classification |
| **PM2_Supporting** | Total allele frequency <0.01% across all genetic ancestry populations in the reference population dataset | Supporting evidence of pathogenic classification |

To enable structured comparison between population criterion fulfillment and ClinVar annotations, concordance was assessed at the variant level by comparing each variant’s ACMG/AMP criteria profile with its corresponding ClinVar pathogenicity classification. Results were stratified by ClinVar review status, which reflects the level of evidentiary support for each classification and is represented on a 0-4 star scale [14]. No globin gene variants have a 3– or 4-star review status. Stratification by ClinVar review status enabled assessment of concordance between ACMG/AMP population criterion fulfillment and curated variant interpretations according to the strength of underlying evidence. These analyses were comparative and were not intended to constitute comprehensive variant reinterpretation or reclassification.

Further, ancestry-exclusive variants were identified to quantify the extent to which observed variation was confined to a single genetic ancestry. An ancestry-exclusive variant was defined as one observed in exactly one gnomAD ancestry group (≥1 alternate allele) and absent from all others. Given substantial sample size differences across genetic ancestries in gnomAD, exclusivity, particularly for rare variants, was interpreted descriptively rather than as evidence of ancestry-specific enrichment. Variant distributions along each gene were examined by quality status, ancestry exclusivity, and fulfillment of the ACMG/AMP criteria defined in Table 1. All data processing, analyses, and visualizations were conducted in R (version 4.4.3) [17].

## Results

### Variant discovery and quality assessment across releases

Across evaluated globin genes, variants in the exome and genome callsets increased from 3,173 in gnomAD v2.1.1 to 13,659 in v4.1, a ∼4-fold increase in variant discovery (Fig.1A). Despite this increase, the proportion of quality-passing variants decreased from 76.6% (2,429/3,173) in v2.1.1 to 47.7% (6,516/13,659) in v4.1 (Fig.1A). In v2.1.1, exomes had a higher pass rate than genomes (80.3% vs 71.0%), which was reversed for v4.1 (41.0% vs 63.2%) (Fig.1A). The v4.1 joint callset had a quality-pass rate of 43.6% (5,021/11,525) (Fig.1A). Of all variants, 2,973 were shared, 200 were unique to v2.1.1, and 10,686 to v4.1 (Fig.1B). Of the 200 variants unique to v2.1.1, 146 passed quality assessment. Nearly all met PM2_Supporting; one met BS1 and one met no population criteria (data not shown). Thus, variants absent from v4.1 rarely provided benign evidence. Among shared variants, quality assessments were mostly concordant (pass-pass 64.2%; fail-fail 15.3%), but discordance was skewed toward pass-to-fail transitions (12.5%) over fail-to-pass transitions (7.9%) (Fig.1B). Across variants carried into the joint callset, 4.4% received a different quality assessment than in the corresponding individual callset (data not shown). Joint calling also identified 56 quality-passing variants absent from both individual callsets (Fig.1C). Accordingly, the quality-passing variant set used for population criterion assessment varies by release and callset. Therefore, release-specific differences in variant detection and quality assessment directly influence BA1, BS1, and PM2_Supporting.

**Figure 1.**
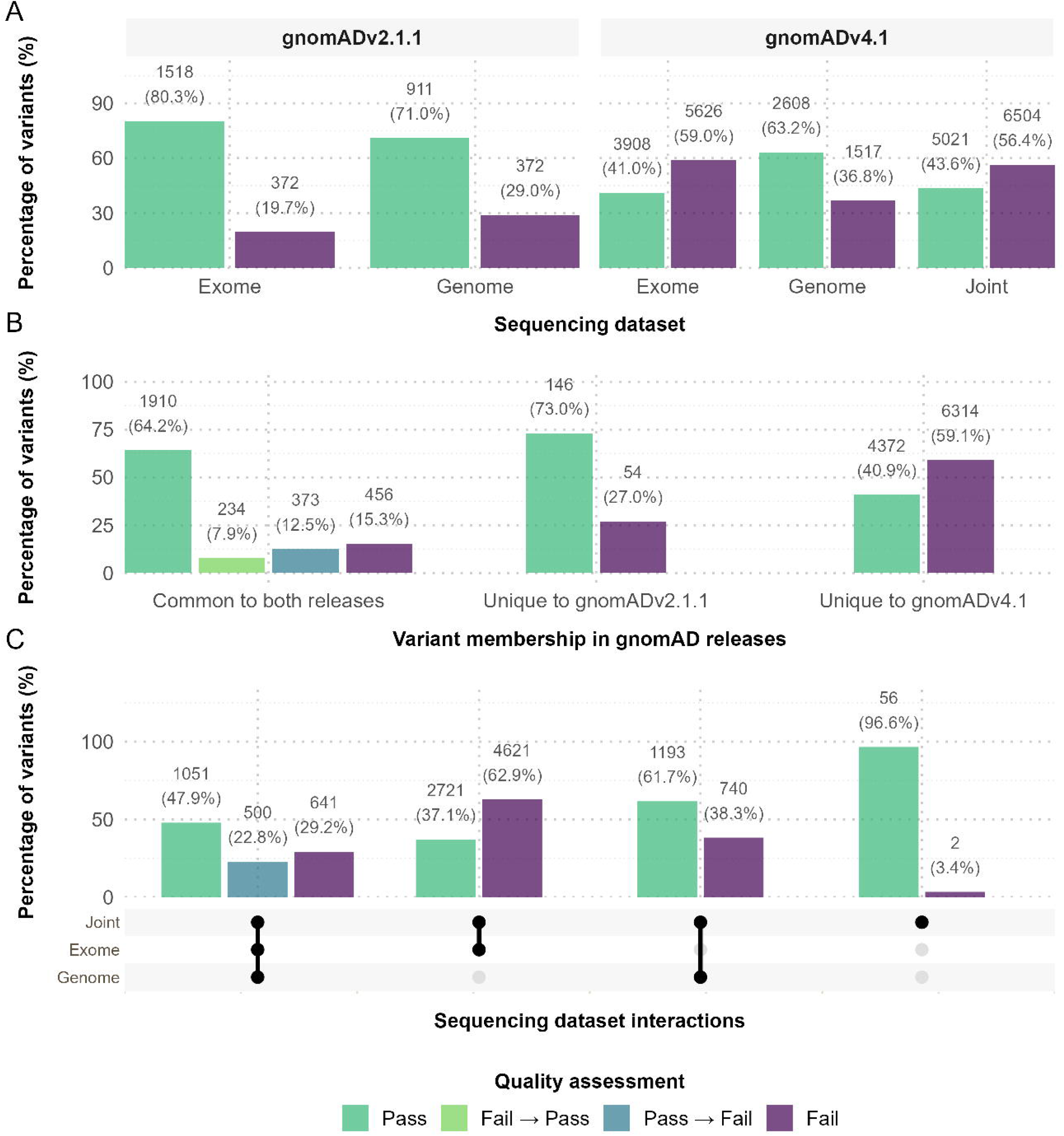
Representation of globin gene variants across gnomAD releases and sequencing datasets. (A) Variant composition of gnomAD v2.1.1 and v4.1 by dataset (exome, genome, joint); green indicates variants that pass quality assessment, and purple indicates quality-failing variants. Percentages denote each dataset’s contribution within a release. (B) Variant membership across releases (exome and genome): shared variants and those unique to v2.1.1 or v4.1. Shared variants are categorized by concordant assessments (pass-pass: green, fail-fail: purple) or assessment changes (pass→fail: blue, fail→pass: light green); percentages are within membership class. (C) Intersections among exome, genome, and joint datasets in gnomAD v4.1. Colors indicate quality-assessment concordance or discordance across callsets: pass (green) and fail (purple) denote concordant assessments, whereas pass→fail (blue) denotes variants that pass in the exome and/or genome callset but fail in the joint callset. No variants were observed failing in either the exome and genome callsets but passing in the joint callset. Percentages denote quality assessment outcomes within each dataset-overlap category.

### Gene-level and ancestry-specific variant distribution

Within the v4.1 joint callset, quality-passing variant counts varied by gene, with the highest counts in *HBB* (1,051; 20.9%), *HBD* (893; 17.8%), and *HBE1* (879; 17.5%) and the lowest in *HBA1* (316; 6.3%) and *HBA2* (359; 7.1%) (Fig.2A). Spatially, quality-passing variants were broadly distributed along *HBB*, *HBD*, and *HBE1*, but clustered mid-gene in *HBG1* and *HBG2*, and toward the 3′ ends of *HBA1* and *HBA2*. For *HBZ*, the quality-passing signal was sparse and mainly confined to intronic sequence (Supplementary Fig. 1). These spatial patterns likely reflect differences in callability rather than functional relevance, particularly in paralogs such as *HBA1*/*HBA2* and *HBG1*/*HBG2*, in which high sequence similarity complicates short-read mapping and gene-specific variant assignment.

**Figure 2.**
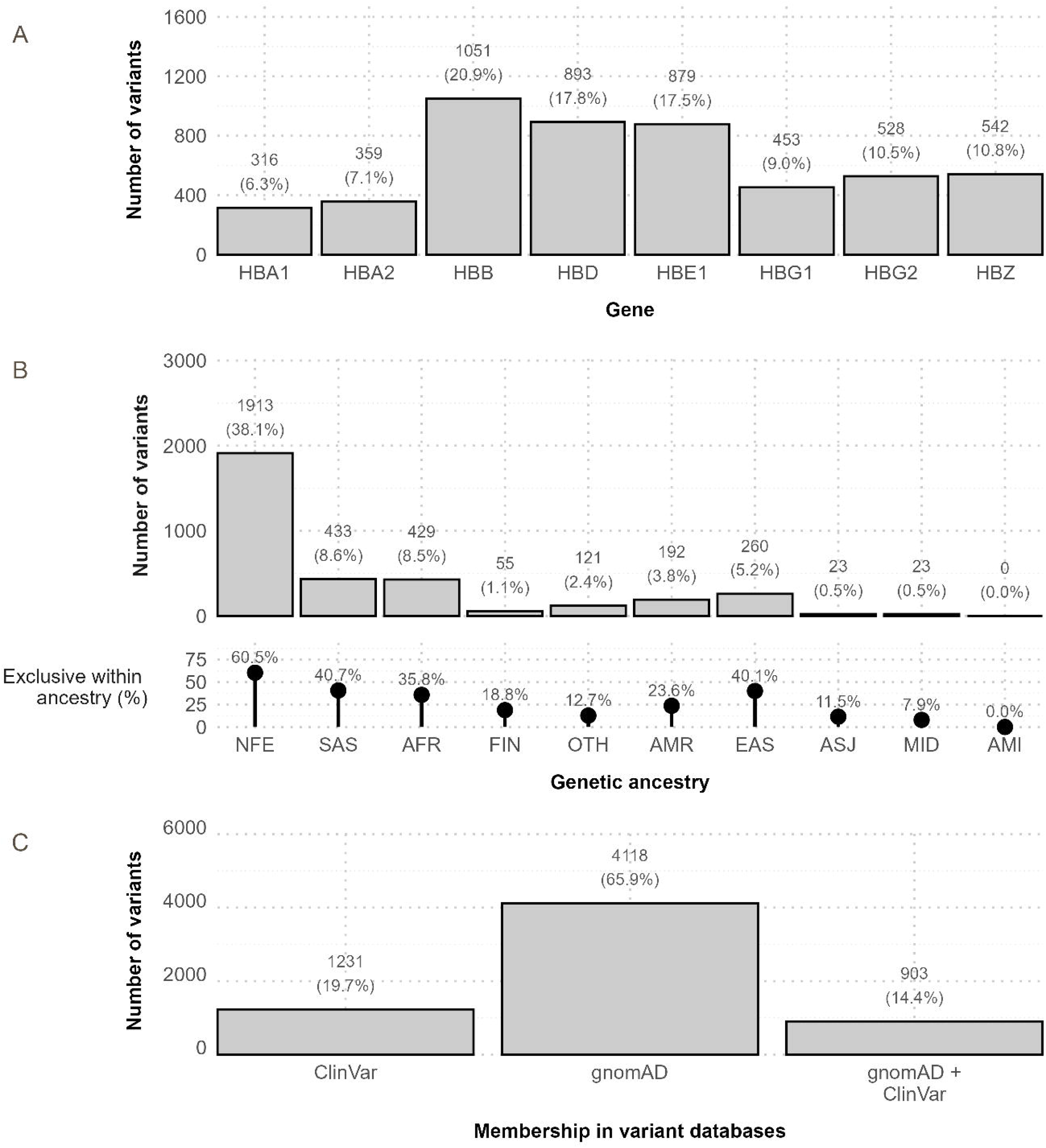
Globin gene variants in gnomAD v4.1 joint dataset that pass quality assessment. (A) Variant counts by globin gene; labels show counts and percentage of the total variant set (n = 5,021). (B) Variants observed exclusively for each genetic ancestry. Bars show, for each genetic ancestry group, the number of variants observed only in that ancestry group; labels give counts and the percentage within the full variant set (n = 5,021). Lollipop chart values represent the exclusivity rate within each ancestry, defined as the proportion of variants observed in that ancestry that are ancestry-exclusive (denominator varies by ancestry). Ancestry groups are presented in descending order based on the number of individuals tested in gnomAD and include: non-Finnish European (NFE), South Asian (SAS), African/African American (AFR), Finnish European (FIN), Other (OTH), Admixed American (AMR), East Asian (EAS), Ashkenazi Jewish (ASJ), Middle Eastern (MID), and Amish (AMI). (C) Database membership across gnomAD and ClinVar: variants present only in ClinVar, only in gnomAD, or in both; labels show counts and percentages indicating each category’s contribution to the total set of variants captured by any of the two databases.

In the v4.1 joint callset, 68.7% of quality-passing variants were observed in only one genetic ancestry; non-Finnish Europeans contributing the largest share overall (1,913/5,021; 38.1%) and within ancestry (1,913/3,162; 60.5%) (Fig.2B). Among other ancestries, within-ancestry exclusivity varied independently of gnomAD sample size ordering; East Asian had 40.1% (260/648) ancestry-exclusive variants, comparable to South Asian (40.7%; 433/1,063) and higher than African/African American (35.8%; 429/1,197) and Finnish (18.8%; 55/293) (Fig.2B). As BA1 and BS1 are evaluated using PopMax AF, typically interpreted across continental (non-founder) populations, the high proportion of ancestry-exclusive variants underscores the importance of increased ancestry representation for robust criterion assessment. Genomic distributions of ancestry-exclusive variants were gene-specific (Supplementary Fig. 2). *HBB*, *HBD*, and *HBE1* showed broadly distributed ancestry-exclusive variants with recurrent local density peaks across multiple ancestries, whereas *HBA1* and *HBA2* were enriched toward the 3′ regions (Supplementary Fig. 2). *HBZ* showed a sparse central signal with clustering toward both gene ends, *HBG1* showed 3′ enrichment with recurrence across ancestries, and *HBG2* showed a denser mid-gene signal with less recurrent overlap (Supplementary Fig. 2). Regions of higher variant density were contributed mainly by larger genetic ancestries, consistent with gnomAD ancestry sample size differences [13].

### Gene-specific constraint metrics across releases

Constraint metrics varied across releases in magnitude and, for some metrics, in the relative rank order of genes (Fig. 3). For LoF intolerance, LOEUF values exceeded 1 for most genes in both releases, indicating tolerance to LoF variation; however, the γ-globin paralogs diverged, with LOEUF decreasing in *HBG2* (0.83 to 0.38) and increasing in *HBG1* (1.13 to 1.64). Using a LOEUF threshold of < 0.6, only *HBG2* was LoF constrained in v4.1 (Fig. 3A). No gene surpassed the missense constraint significance cutoff (z≥3.09) in either release, indicating missense variation tolerance; however z-scores shifted in magnitude and rank: the highest value shifted from *HBA1* in v2.1.1 (1.53; 0.80 in v4.1) to *HBG2* in v4.1 (2.22; 1.35 in v2.1.1), *HBB* changed sign (–0.21 to 0.78), and *HBD* remained negative (–1.01 to –0.20) (Fig.3B) [13]. Synonymous constraint z-scores also shifted toward a narrower range in v4.1, attenuating extreme values relative to v2.1.1, including decreases at previously elevated genes (*HBA1*: 2.03 to 0.04; *HBG1*: 2.10 to 0.40) and movement toward zero at the most negative gene (*HBB* –3.80 to –1.41), consistent with improved constraint estimation in v4.1 (Fig.3C).

**Figure 3.**
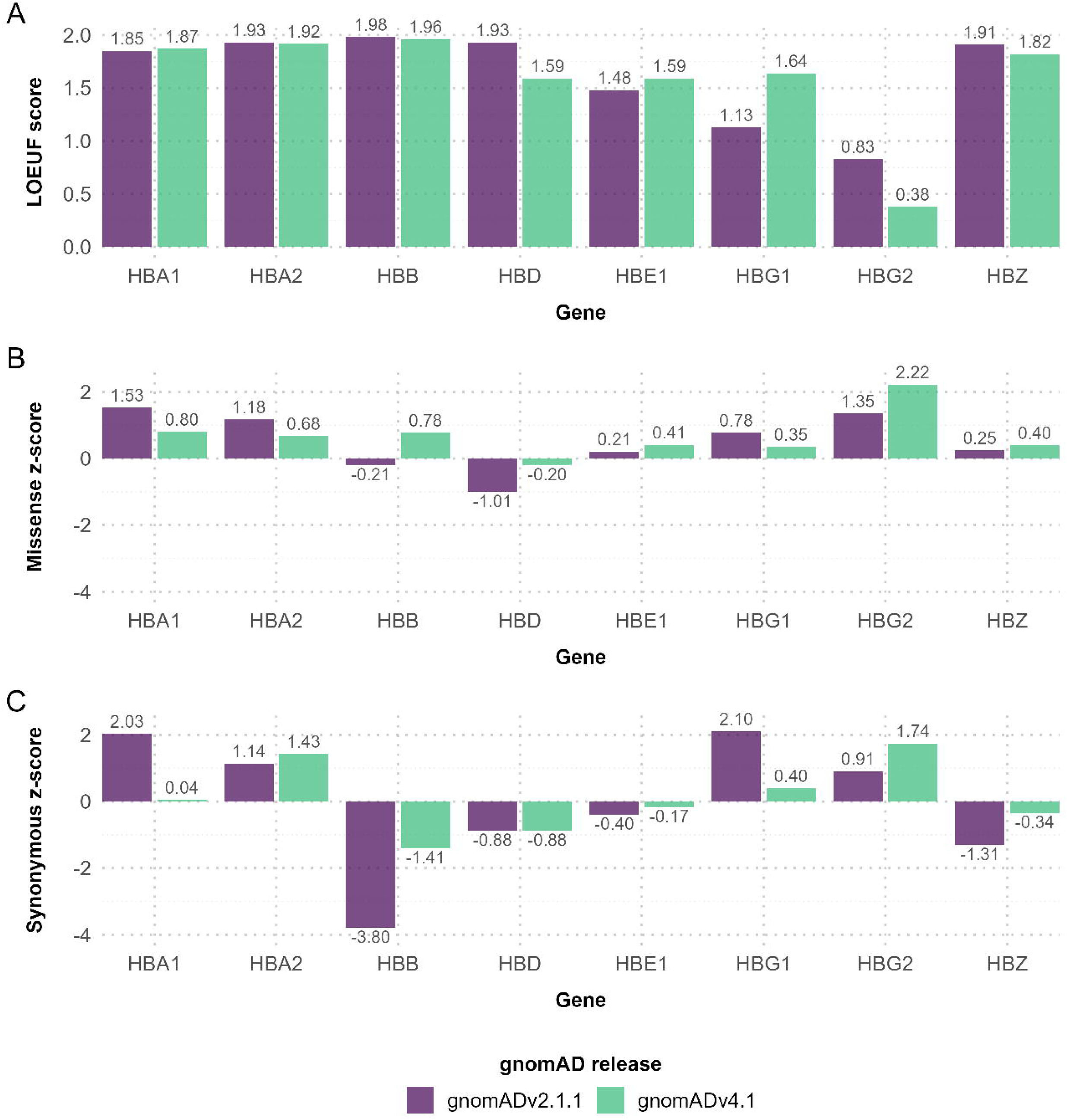
Gene-level constraint across globin genes in gnomAD. Constraint metrics for *HBA1*, *HBA2*, *HBB*, *HBD*, *HBE1*, *HBG1*, *HBG2*, and *HBZ* are shown for gnomAD v2.1.1 (purple) and v4.1 (green). (A) Loss-of-function (LoF) intolerance is reported as the LoF observed/expected upper bound fraction (LOEUF), with lower values indicating stronger depletion of LoF variants relative to expectation. (B) Missense constraint is shown as a z-score. (C) Synonymous constraint is shown as a z-score. For (B) and (C), larger positive values for constraint z-scores indicate stronger depletion of observed variants relative to expectation, whereas increasingly negative z-scores indicate enrichment of observed variants relative to expectation.

### Impact on ACMG/AMP population criterion fulfillment

Across releases, PM2_Supporting was the most commonly met population criterion among quality-passing variants, increasing 2.6-fold from 1,781 (81.5%) in v2.1.1 to 4,695 (93.5%) in v4.1, with 2,914 additional variants meeting PM2_Supporting (Fig.4A). The increase in PM2_Supporting reflected newly detected rare variants in v4.1 rather than reclassification of shared variants, consistent with the larger v4.1 sampling framework. In contrast, the proportion of variants meeting benign evidence criteria based on elevated PopMax AF or homozygote observations was lower in v4.1 than in v2.1.1 (Fig. 4A). Fulfillment of conflicting population criterion combinations was infrequent (Supplementary Table 1); nonetheless, PM2_Supporting co-occurred with BS1 in 0 variants in v2.1.1 and 9 (0.2%) in v4.1 and co-occurred with BS2 in 3 (0.1%) and 32 (0.6%), respectively (Fig.4B). Within v4.1, BS1/PM2_Supporting co-occurrences were fewer in the joint callset than in the individual callsets considered together (9 vs 20), whereas BS2/PM2_Supporting co-occurrences were modestly higher (32 vs 27) (data not shown). This pattern is consistent with more stable allele frequency estimation in the joint callset, reducing BS1/PM2_Supporting conflicts driven by borderline frequency estimates while strengthening confidence in BS2 assignments for rare variants observed in the gnomAD reference population. Among shared variants, changes involving PM2_Supporting were unevenly distributed, with more variants meeting PM2_Supporting in v4.1 (58 variants; 3.5%) than no longer meeting it (24 variants; 1.5%) (Fig.4C). As extremes of variant fulfillment across criteria, PM2_Supporting fulfillment was high and ranged from 82.3% in *HBG1* to 98.9% in *HBA2*, whereas BA1 and BS1 were particularly rare in *HBA1* and *HBA2* (≤0.9%) (Fig.5A). *HBG1* showed the highest proportions of BA1 (9.1%) and BS2 (13.9%), while *HBA2* showed the highest proportion of BS2_Supporting (8.1%) (Fig.5A). PM2_Supporting variants were broadly distributed along genes, whereas BA1, BS1, BS2, and BS2_Supporting variants were sparse (Supplementary Fig. 3).

**Figure 4.**
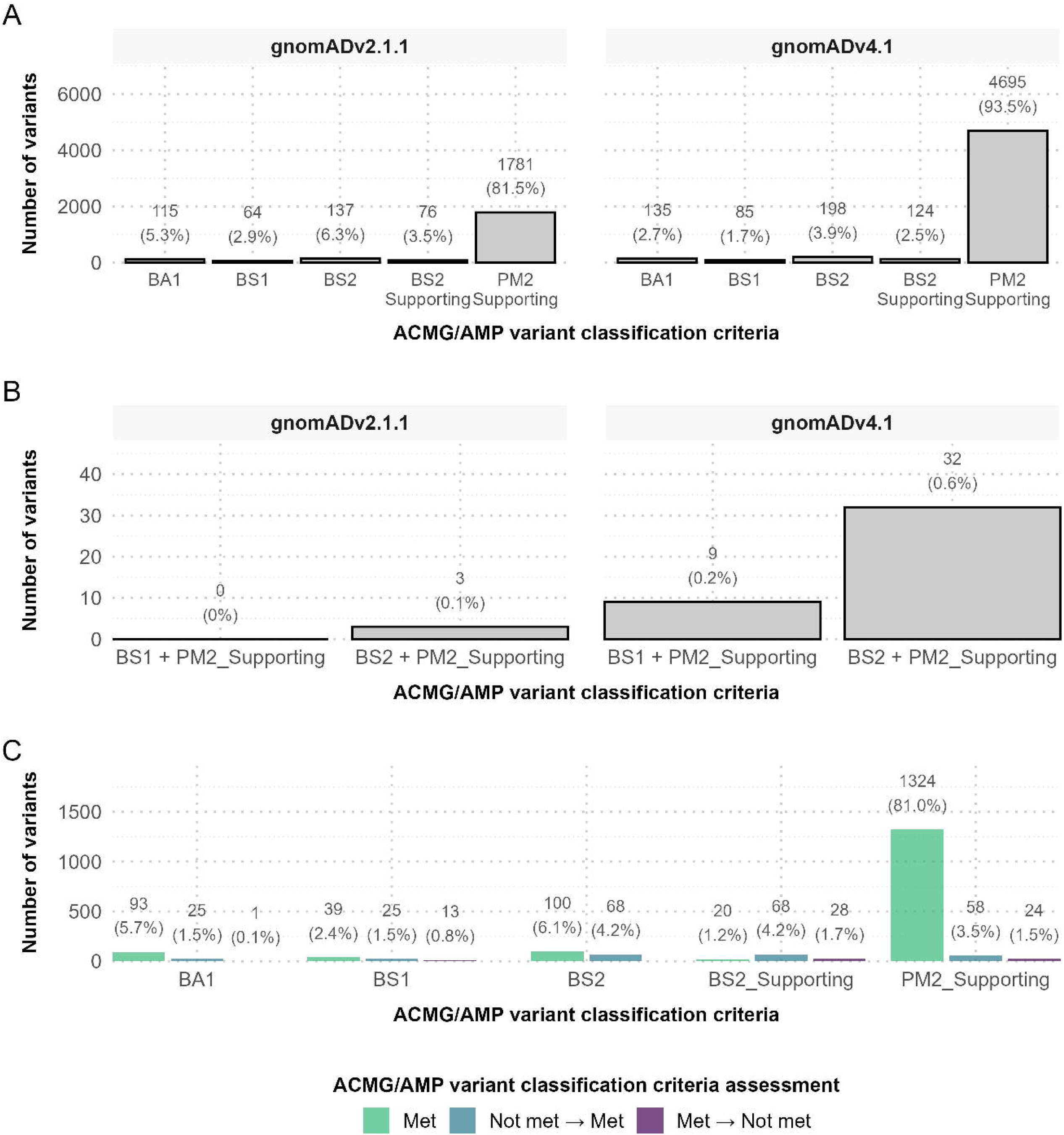
ACMG/AMP criterion assessment for globin gene variants in gnomAD. (A) Counts of quality-passing variants meeting BA1, BS1, BS2, BS2_Supporting, or PM2_Supporting in gnomAD v2.1.1 and v4.1; percentages are within release. (B) Subset of panel A meeting criterion combinations supporting conflicting evidence (BS1 with PM2_Supporting; BS2 with PM2_Supporting). (C) For quality-passing variants present in both releases, criterion status across releases: met in both (green), not met in v2.1.1 but met in v4.1 (blue), or met in v2.1.1 but not met in v4.1 (purple); percentages are calculated within each criterion.

**Figure 5.**
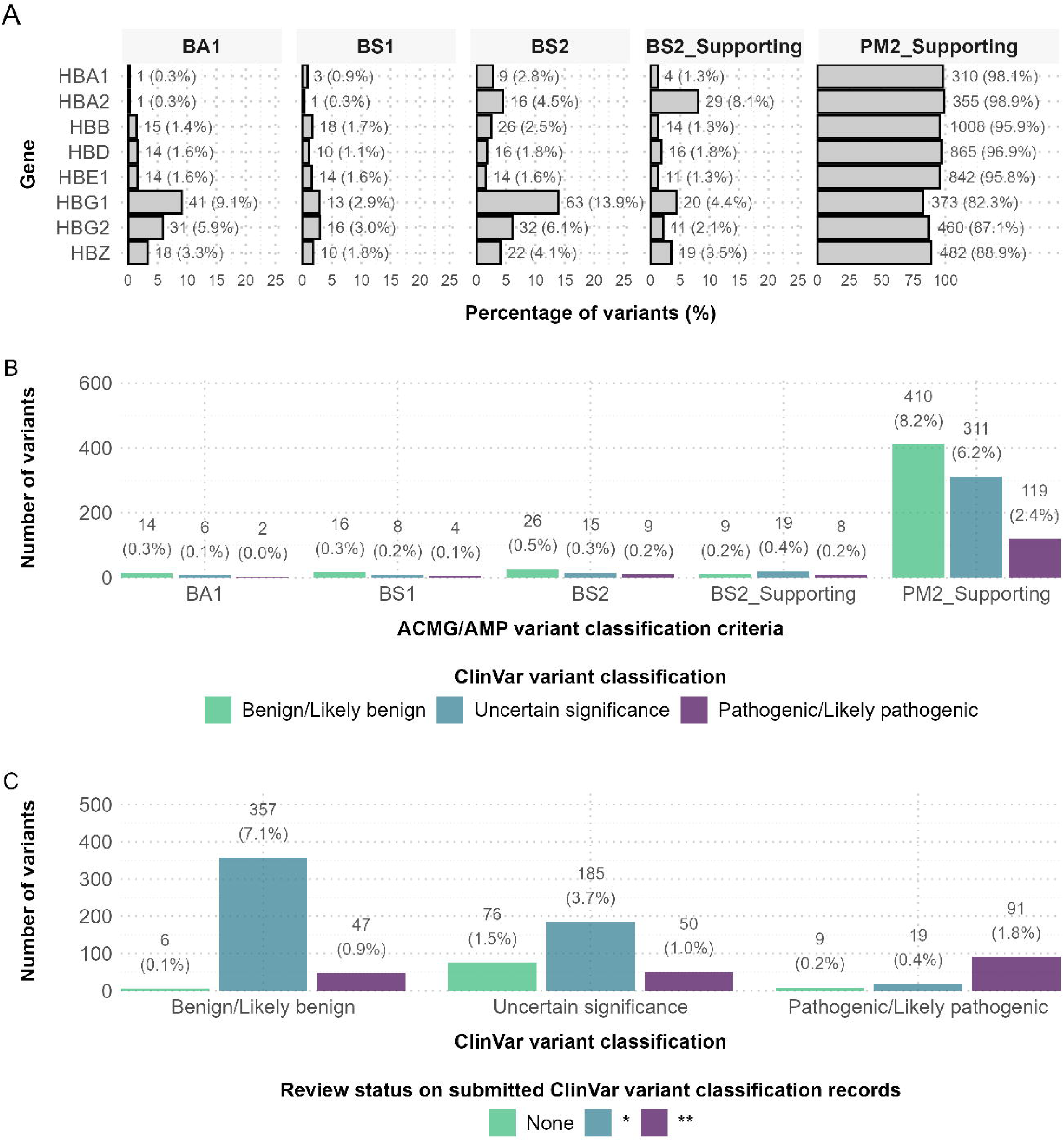
ACMG/AMP criteria and ClinVar annotations for globin gene variants. All panels summarize variants from the gnomAD v4.1 joint dataset that pass quality assessment. (A) Percentage of variants meeting BA1, BS1, BS2, BS2_Supporting, or PM2_Supporting, stratified by gene; percentages are within gene. (B) Number of variants present in ClinVar, grouped by criterion and ClinVar clinical significance (benign/likely benign: green, uncertain significance: blue, pathogenic/likely pathogenic: purple); percentages are relative to all 5,021 gnomAD v4.1-represented variants. (C) ClinVar-annotated PM2_Supporting variants stratified by clinical significance and ClinVar review status (none: green, ★: blue, ★★: purple); percentages are within clinical-significance class, relative to all 5,021 gnomAD v4.1-represented variants.

### Overlap with clinical annotation resources

ClinVar overlap was limited, with 903 of 5,021 quality-passing gnomAD variants also reported in ClinVar, 4,118 observed only in gnomAD v4.1, and 1,231 reported only in ClinVar (Fig.2C). Among variants overlapping ClinVar, those meeting BA1, BS1, BS2, or BS2_Supporting were mostly classified as benign or likely benign, although a small number carried uncertain significance (48) or pathogenic/likely pathogenic (23) assertions, whereas variants meeting PM2_Supporting showed a broader distribution across benign/likely benign (410), uncertain significance (311), and pathogenic/likely pathogenic (119) classifications (Fig.5B). For PM2_Supporting variants with ClinVar records, pathogenic or likely pathogenic interpretations were more often supported by multiple-submitter, non-conflicting submissions (two stars; 91/119; 76.5%) than benign or likely benign interpretations (47/410; 11.5%), which were predominantly single-submitter or unassessed submissions (Fig.5C).

## Discussion

This study compares globin gene variant representation in gnomAD v2.1.1 and v4.1 and evaluates how ClinGen Hemoglobinopathy VCEP adoption of v4.1 may affect the availability and strength of population evidence for globin gene variant interpretation, with implications for other gene-disease contexts. The transition to gnomAD v4.1 represents a step-change in sampling scale and ancestry composition, with updated variant calling, filtering, and frequency estimation [13]. Differences between v2.1.1 and v4.1 therefore reflect cohort enlargement and release-specific analytical models, not simple variant accrual [13].

A substantial increase in observed globin gene variants accompanied this transition, although the percentage of quality-passing variants decreased in v4.1, and newly detected variants were more likely to fail quality assessment than variants present in both releases. Quality assessment for variants shared across releases was generally stable, but discordance was present and more often reflected pass-to-fail than fail-to-pass transitions. Release sensitivity has also been reported in movement disorders and cystic fibrosis, in which variants absent from v2.1.1 appear in v4.1 and revised allele-frequency estimates affect variant interpretation [12,18,19]. The present study extends prior work by systematically assessing ACMG/AMP population criteria fulfillment across all variants, rather than limiting analysis to case-level reinterpretation or curated disease-associated variant sets. Further, joint calling increased variant discovery by identifying variants absent from either individual callset, supporting the joint callset as the most comprehensive resource for variant interpretation [20]. Because BA1, BS1, BS2, BS2_Supporting, and PM2_Supporting use allele frequencies and homozygote counts from quality-filtered callsets, population evidence and criterion fulfillment are sensitive to both gnomAD release and callset. This sensitivity is most consequential for variants near frequency and homozygote thresholds, for which small shifts in allele counts or quality assessment can change whether population evidence is gained or lost for the same variant across releases. Routine reporting of gnomAD release and callset is therefore essential, especially for borderline variants [16,18].

Differences in quality-passing variant counts across globin genes were most consistent with differences in callable sequence length and regional callability. *HBB* had the most quality-passing variants, consistent with a larger callable sequence and more reliable calling than the highly sequence-similar α– and γ-globin gene paralogs, which yielded fewer quality-passing variants in regions of reduced mappability and alignment ambiguity [21–23]. This also likely reflects gnomAD’s reliance on short-read sequencing, which reduces accuracy in paralogous and repetitive regions that long-read sequencing may better resolve [13]. The genomic distribution of pass and fail variants supports this interpretation, with quality-failing variants clustering in discrete intervals that shrink the callable portion of globin genes. HBZ illustrates this clearly, with a prominent cluster of quality-failing variants near the repetitive, GC-rich α-globin locus control region [24].

Augmented sampling and broader ancestry representation in gnomAD v4.1 increased detection of rare globin gene variants, many of which were observed in only a single genetic ancestry. Such apparent ancestry exclusivity is expected under unequal sampling as the probability of detecting an extremely rare variant rises with the number of alleles sampled within a genetic ancestry, while the same variant may remain undetected in smaller genetic ancestries due to limited sampling depth [13,25]. Accordingly, rare variant exclusivity should be interpreted as contingent on current sampling, rather than true absence from other populations [25]. However, within-ancestry exclusivity did not strictly follow gnomAD sample size ordering, indicating that sample size alone does not fully explain the observed patterns. These ancestry sampling differences directly affect ACMG/AMP frequency criteria. Because BA1 and BS1 rely on ancestry-stratified PopMax AF, variation in sample size and ancestry representation can shift benign criterion fulfillment [7]. Broader representation yields more reliable frequency estimates, whereas underrepresented genetic ancestries remain more susceptible to imprecise estimates near VCEP thresholds [25].

Changes in gene-level constraint metrics between gnomAD releases underscore how increased sample size and updated analytical methods can influence interpretation. Synonymous constraint z-scores are expected to be near neutral at the gene-level but may be distorted in paralog-rich genes such as *HBA1*/*HBA2* and *HBG1*/*HBG2* by technical artefacts [13,16]. *HBB* retained a negative synonymous constraint z-score in both releases, but its reduced magnitude in v4.1 likely reflects improved data quality and recalibrated expectation models rather than true biological constraint. The concurrent movement of *HBA1* and *HBG1* synonymous constraint z-scores toward neutrality supports a shared technical refinement explanation [13]. Beyond synonymous constraint, most globin genes remained tolerant to LoF and missense variation across releases, despite shifts in constraint estimates’ magnitude and direction. This pattern is consistent with the predominantly recessive inheritance of hemoglobinopathies, in which deleterious alleles can persist in clinically unaffected heterozygous individuals [3,26,27]. *HBG2* was the only gene with a LOEUF score consistent with LoF constraint in v4.1, indicating that LoF depletion is heterogeneous even among paralogs. This is biologically plausible given non-identical regulation and expression dynamics during fetal development, during which disruption of one paralog, particularly *HBG2*, may have a greater effect on total fetal hemoglobin production [28,29].

The clinical relevance of gnomAD augmentation is most apparent in ACMG/AMP population criterion fulfillment. PM2_Supporting was met by most quality-passing variants in both releases and was more prevalent in v4.1, consistent with the influx of newly detected rare variants [5]. By contrast, criteria supporting benign classification were met by comparatively few variants and were proportionally less frequent in v4.1 than v2.1.1. These findings indicate that most callable globin gene variants are rare and remain rare despite increased sampling; PM2_Supporting is therefore met for most and has limited discriminative power in isolation. Hemoglobinopathy-causing variants may be observed in gnomAD through the inclusion of unaffected heterozygous individuals, and some may reach higher ancestry-specific frequencies through selection [3]. Therefore, presence in gnomAD alone should not be interpreted as evidence against pathogenicity. Accordingly, the Hemoglobinopathy VCEP has defined, in line with ClinGen recommendations, a list of variants excluded from benign population frequency criteria [5]. Under these conditions, the main effect of gnomAD augmentation is to strengthen population evidence by narrowing uncertainty around allele frequency estimates rather than altering interpretive implications. Interpretive shifts are therefore expected mainly when BA1/BS1 thresholds are crossed, although PopMax AF estimates and threshold attainment remain contingent on ancestry representation in gnomAD [5,7,13,25]. This underscores the need to validate ClinGen Hemoglobinopathy VCEP thresholds following major gnomAD updates.

Variants meeting conflicting population criteria combinations were scarce, and most shared variants retained concordant criterion profiles, indicating ClinGen Hemoglobinopathy VCEP population thresholds remain robust under gnomAD augmentation. However, instances in which PM2_Supporting fulfillment co-occurred with BS2 warrant cautious interpretation, as gnomAD lacks detailed phenotypic information and the potential influence of co-inherited modifying factors in homozygous individuals cannot be excluded. Nonetheless, shifts in criterion fulfillment demonstrate that population evidence remains release– and callset-dependent. This dependence is most consequential for borderline variants, for which modest changes can determine whether sufficient evidence accrues to support a likely classification, especially when cumulative evidence is limited or conflicting [11,18]. In this setting, review by the ClinGen Hemoglobinopathy VCEP is essential to contextualize release-sensitive population evidence within the full ACMG/AMP framework, particularly for genes other than *HBA1*, *HBA2*, and *HBB*, for which thresholds have not been formally specified. Population evidence for curated variants should therefore be reassessed as gnomAD evolves, with explicit reporting of release and callset and prioritization of variants near frequency thresholds.

Most quality-passing globin gene variation observed in gnomAD lacked ClinVar annotation. This reflects scope differences as gnomAD catalogs population variation, whereas ClinVar aggregates clinical interpretation submissions and is enriched for variants encountered and annotated in diagnostic settings [13,14]. Therefore, benign variants are often under-submitted, and many newly observed rare variants in gnomAD may not yet have been clinically evaluated. Conversely, ClinVar may contain variants absent from gnomAD due to ascertainment outside reference cohorts. Limited overlap therefore highlights a translational gap between population data and clinical interpretation resources. Moreover, ClinVar review status indicates that evidentiary support varies across classification categories. Pathogenic assertions more often reflected multiple concordant submissions, whereas benign assertions are more often supported by single-submitter records or submissions with conflicting evidence. Accordingly, variants labelled as benign in ClinVar but supported only by single-submitter or limited evidence may instead be classified as variants of uncertain significance under stricter evidentiary standards. Interpretation is further constrained by the review-status framework itself, as expert-panel assertions are absent for globin genes, capping the maximum review tier available at two stars [14]. This imbalance limits the use of ClinVar alone to assess concordance between population evidence and clinical classification, and underscores the role of the ClinGen Hemoglobinopathy VCEP in providing harmonized, disease-specific interpretations [14].

## Conclusion

In summary, the transition from gnomAD v2.1.1 to v4.1 substantially increased globin gene variant discovery and shifted the observed landscape toward rarer variants, with joint calling extending discovery beyond single-modality callsets. Release– and callset-specific differences in variant filtering and constraint metrics underscore that population evidence is dataset-dependent and should be applied with explicit reporting of the gnomAD release and callset queried. Using ClinGen Hemoglobinopathy VCEP specifications, ACMG/AMP population criterion fulfillment was mostly stable for shared variants, while the influx of newly detected rare variants increased PM2_Supporting prevalence and highlighted its limited discriminative value when considered in isolation. Although benign evidence criteria were less frequently met, variants exceeding BA1/BS1 thresholds or supported by homozygote observations (BS2/BS2_Supporting) were mostly concordant with benign ClinVar assertions, supporting the specificity of calibrated frequency and homozygosity thresholds. Because most gnomAD variants lacked ClinVar annotations, concordance could be evaluated for only a minority of variants, underscoring the need for disease-specific expert curation. Collectively, these findings support the central role of gnomAD v4.1 in hemoglobinopathy variant interpretation, while emphasizing that population evidence is one component of ACMG/AMP classification that must be integrated with clinical, functional, segregation, and case-level evidence.

## Data Availability

gnomAD and ClinVar data were obtained from their respective open-access repositories: gnomAD (https://gnomad.broadinstitute.org/downloads; accessed 28 May 2025) and ClinVar (https://ftp.ncbi.nlm.nih.gov/pub/clinvar/vcf_GRCh38/; accessed 10 October 2025). All analysis scripts are publicly available at https://github.com/2238154/gnomAD-Manuscript.git.

## Author Contributions

M.K., C.S., and P.K. were responsible for study design. M.K. and M.X. developed the methodology. M.K. performed the formal analysis and prepared the visualizations. C.S. and P.K. supervised the work. M.K. wrote the original draft. M.K., M.X., C.S., P.K., and C.W.L. reviewed and edited the manuscript.

## Acknowledgments

This work was co-funded by the EU Horizon 2020 research and innovation programme under grant agreement No 101017549 (GenoMed4All) and by COST Action CA22119: Haemoglobinopathies in European Liaison of Medicine and Science (HELIOS), supported by COST (European Cooperation in Science and Technology). The authors acknowledge the use of ChatGPT 5.4 (accessed February 2026) for grammatical proofreading and to improve the clarity and readability of the manuscript. All AI-assisted output was reviewed, revised, and approved by the authors, who take full responsibility for the accuracy and integrity of the work.

