## Supplementary Figures for "Analyzing globin gene variation in gnomAD: implications for variant interpretation in hemoglobinopathies"

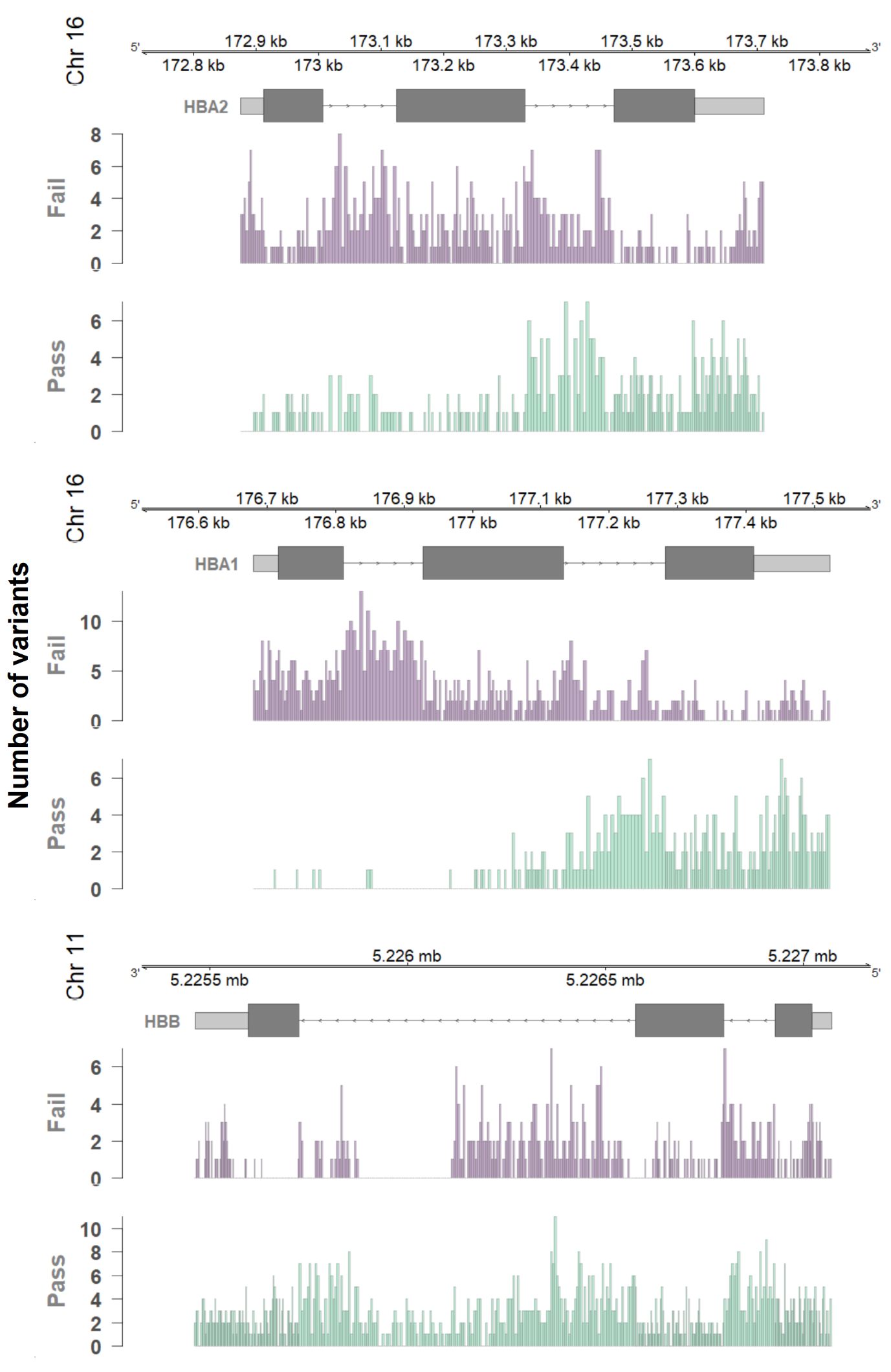

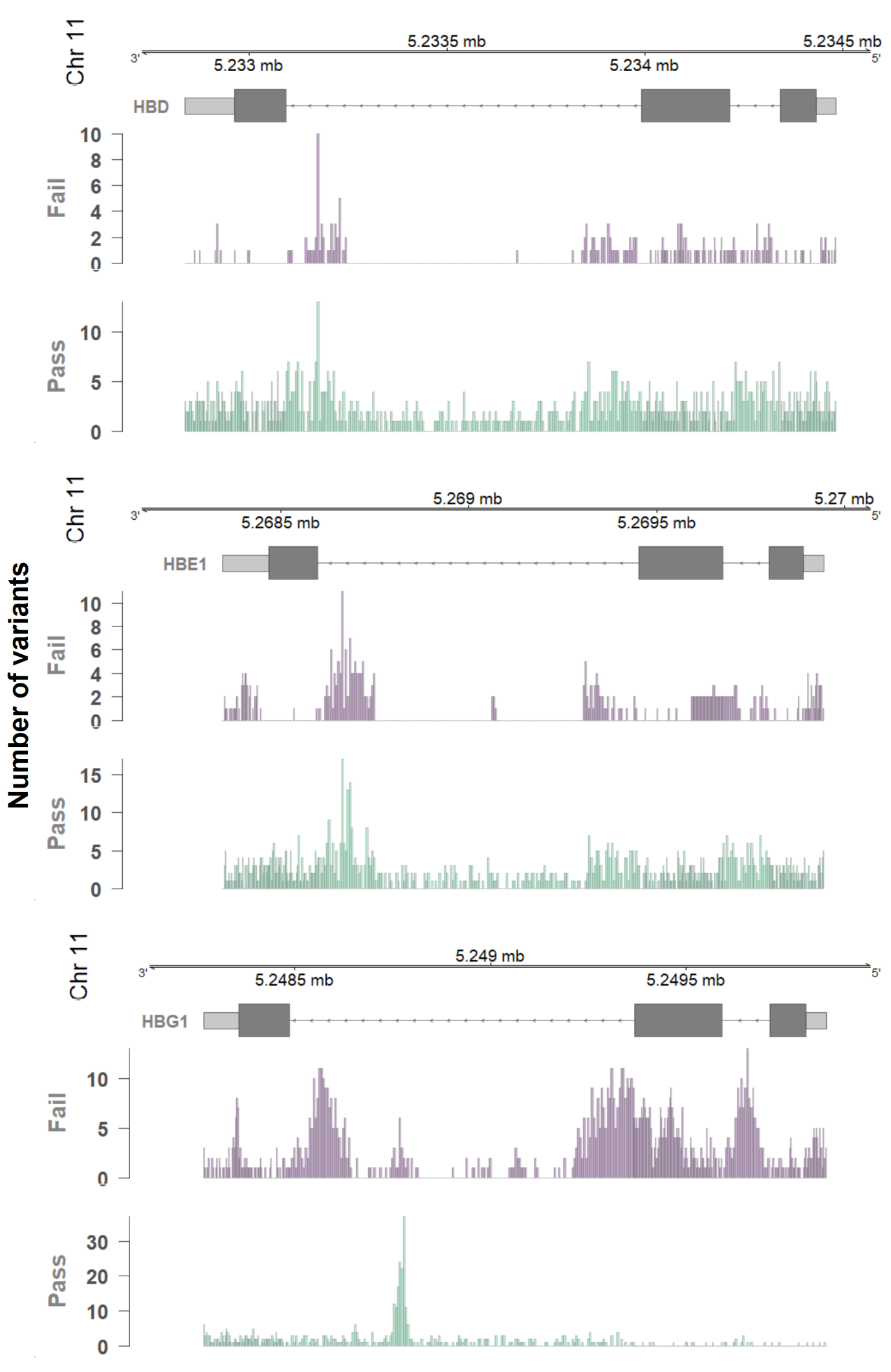

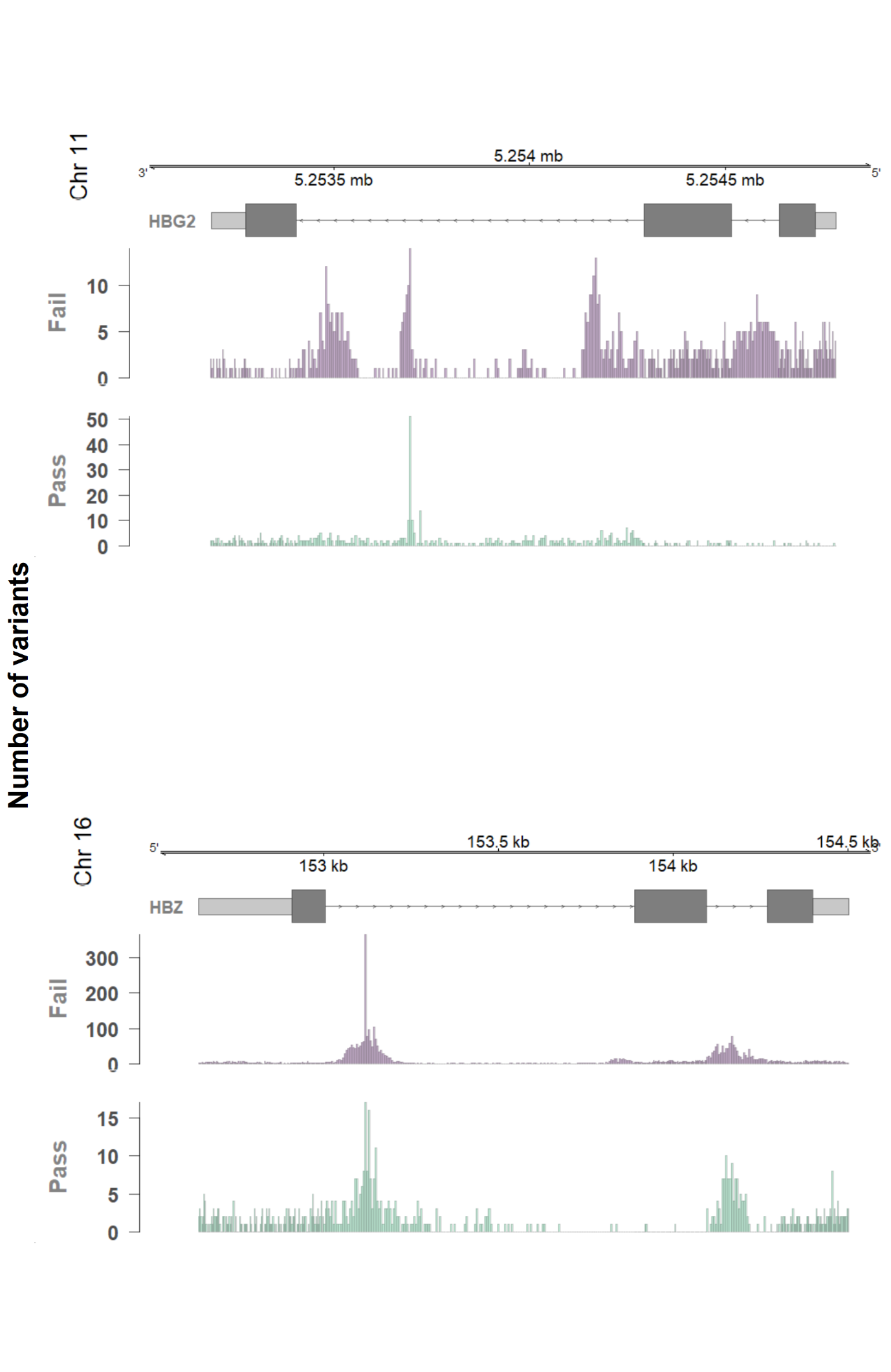

**Supplementary Figure 1. Genomic distribution of globin gene variants in gnomAD v4.1 by quality status.** Histograms show the distribution of variants from the gnomAD v4.1 joint dataset across *HBA2*, *HBA1*, *HBB*, *HBD*, *HBE1*, *HBG1*, *HBG2*, and *HBZ*, stratified by quality assessment (**pass**, green; **fail**, purple). In each track, bar height represents the number of variants per genomic bin, using 3-bp bins in exonic regions and 5-bp bins in intronic regions. Gene models and genomic coordinates are shown above each panel; where horizontal lines denote introns, light grey boxes denote untranslated regions, and dark grey boxes denote coding sequences.

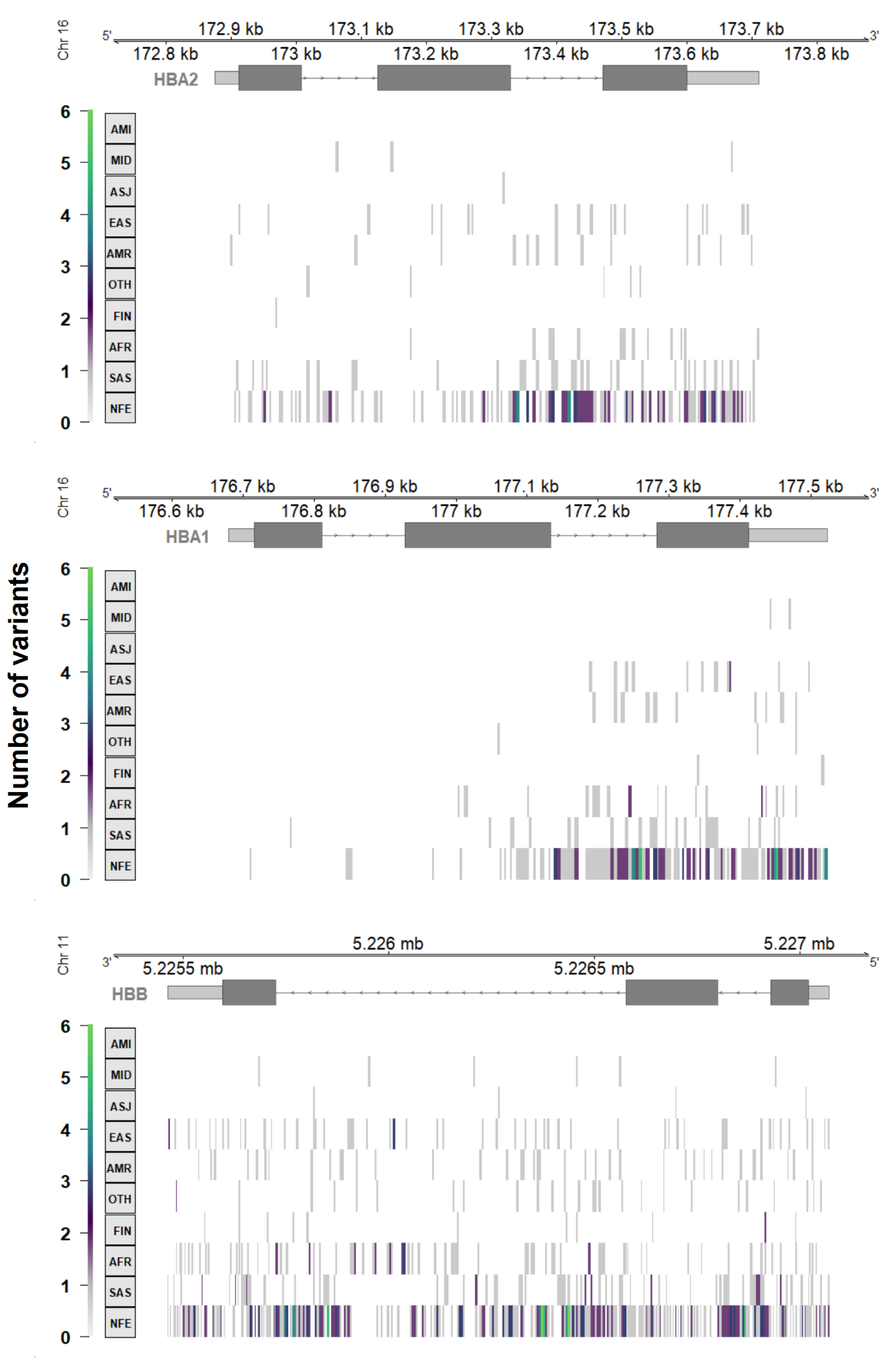

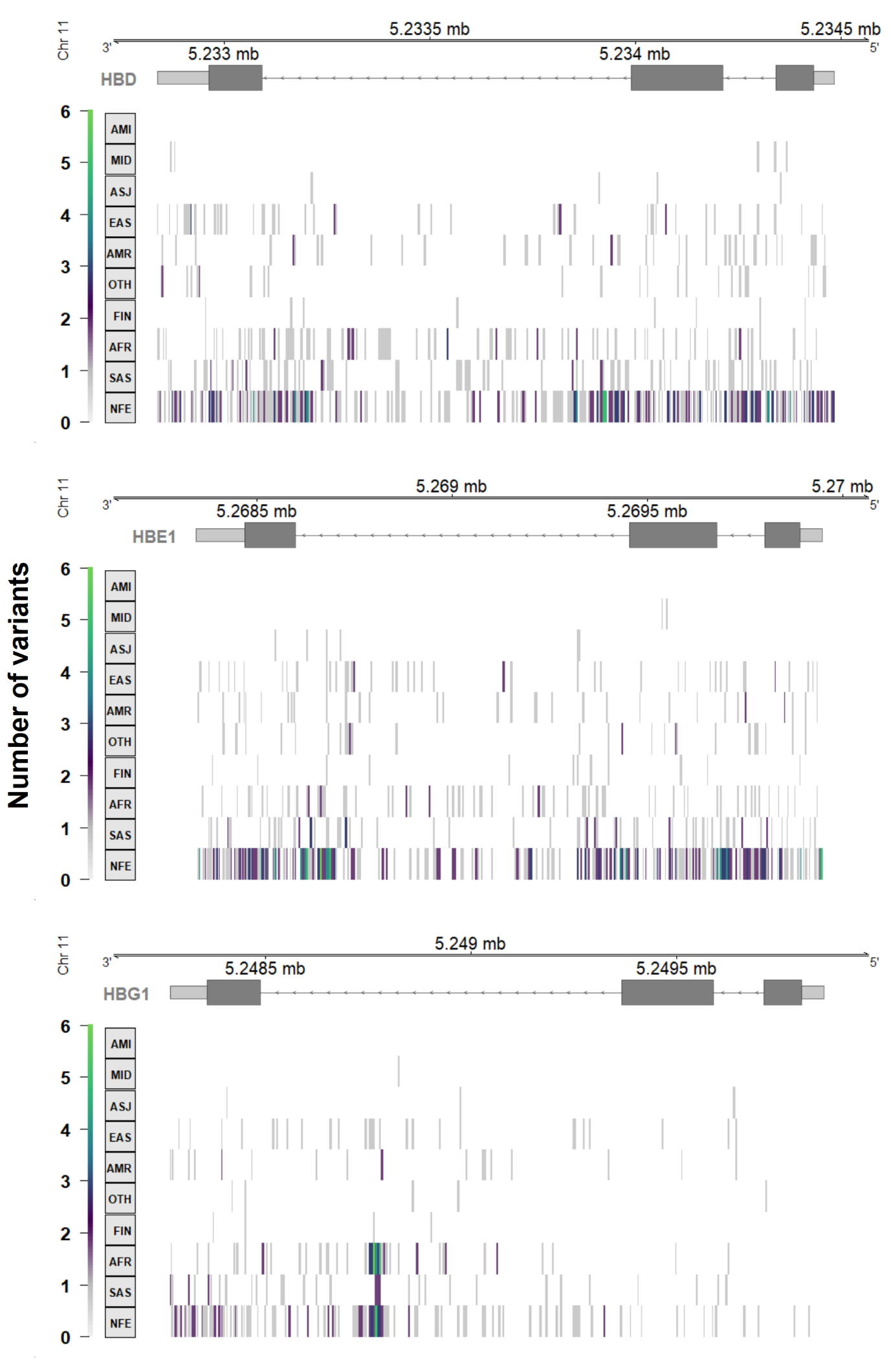

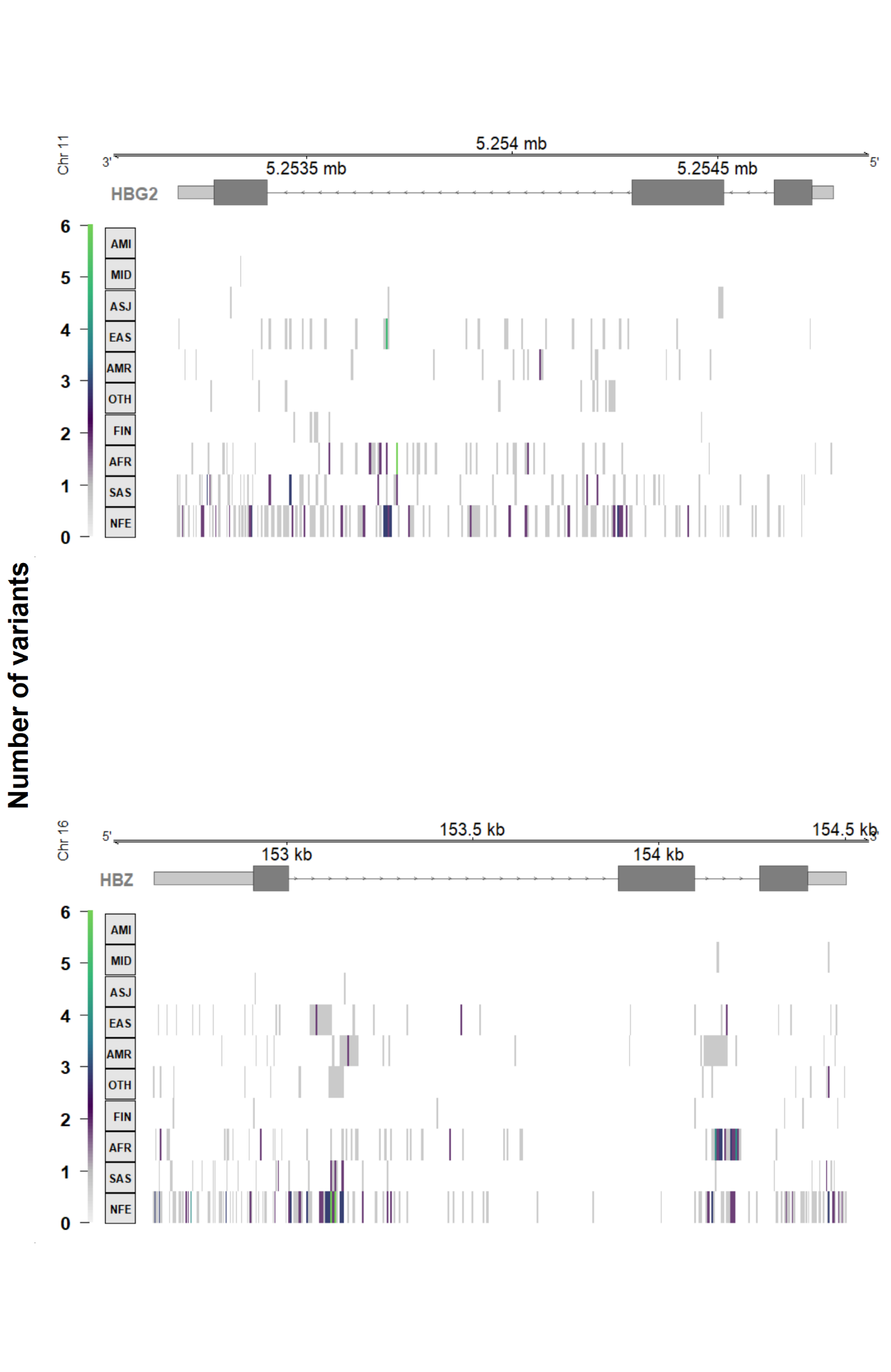

**Supplementary Figure 2. Ancestry-exclusive globin gene variants in gnomAD v4.1.** Heatmaps show the genomic distribution of quality-passing variants from the gnomAD v4.1 joint dataset observed exclusively within a single genetic ancestry across *HBA2*, *HBA1*, *HBB*, *HBD*, *HBE1*, *HBG1*, *HBG2*, and *HBZ*; shading reflects the count of ancestry-exclusive variants per genomic bin. Ancestry groups are presented in ascending order based on the number of individuals tested in gnomAD and include: Amish (AMI), Middle Eastern (MID), Ashkenazi Jewish (ASJ), East Asian (EAS), Admixed American (AMR), Other (OTH), Finnish European (FIN), African/African American (AFR), South Asian (SAS), and non-Finnish European (NFE). Gene models and genomic coordinates are shown above each panel; where horizontal lines denote introns, light grey boxes denote untranslated regions, and dark grey boxes denote coding sequences. Gene tracks are displayed using 3-bp exonic bins and 5-bp intronic bins.

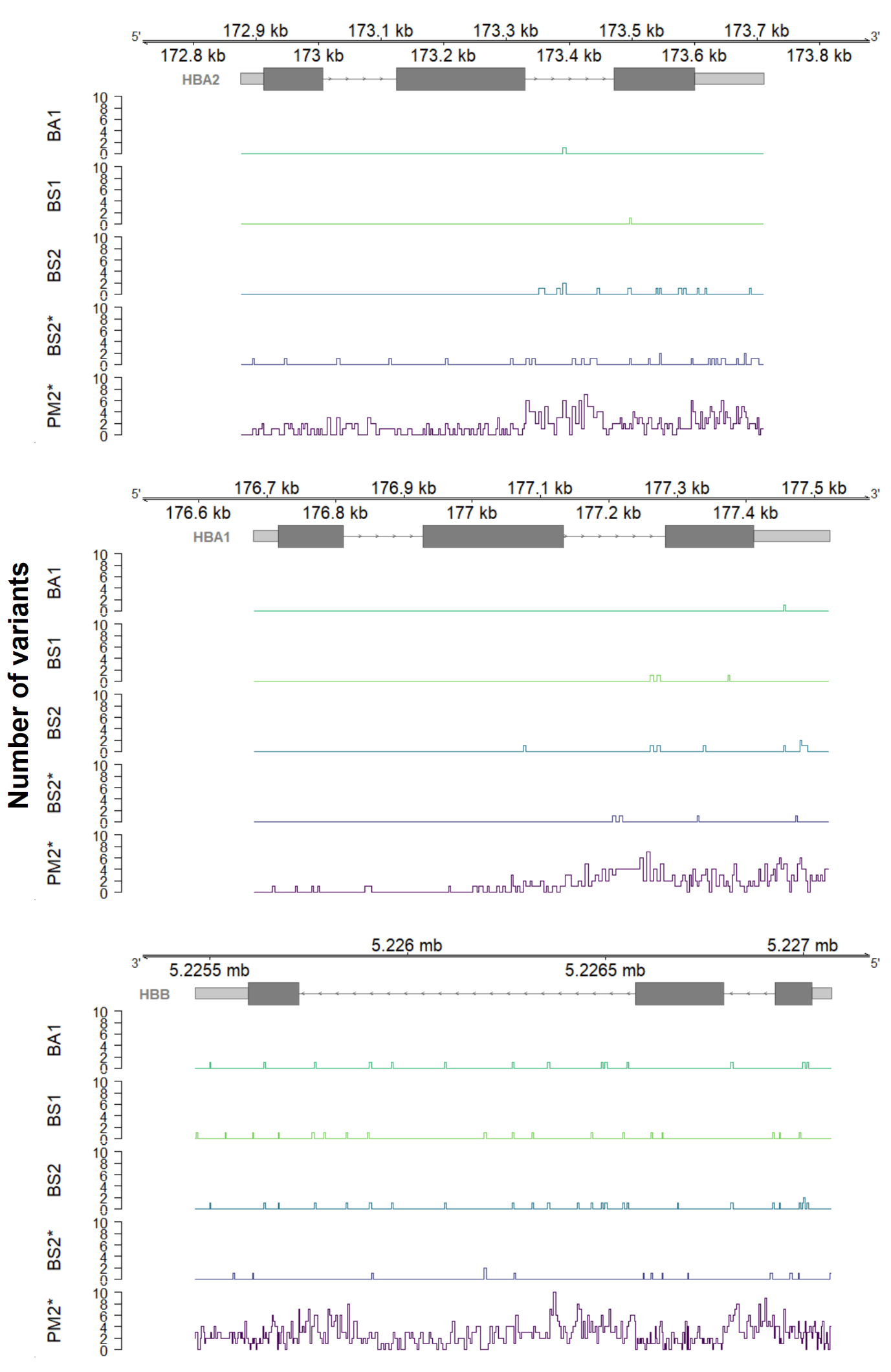

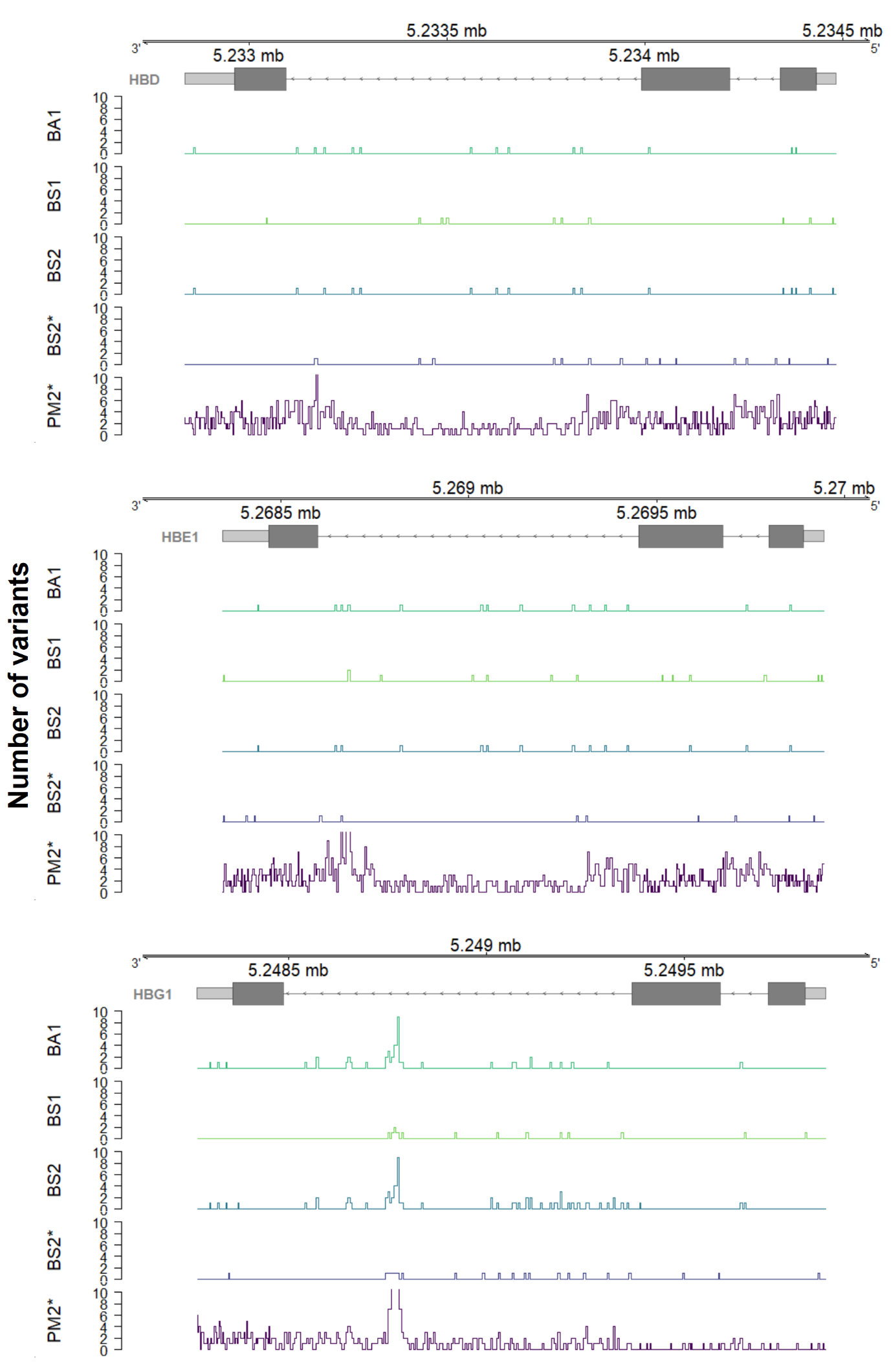

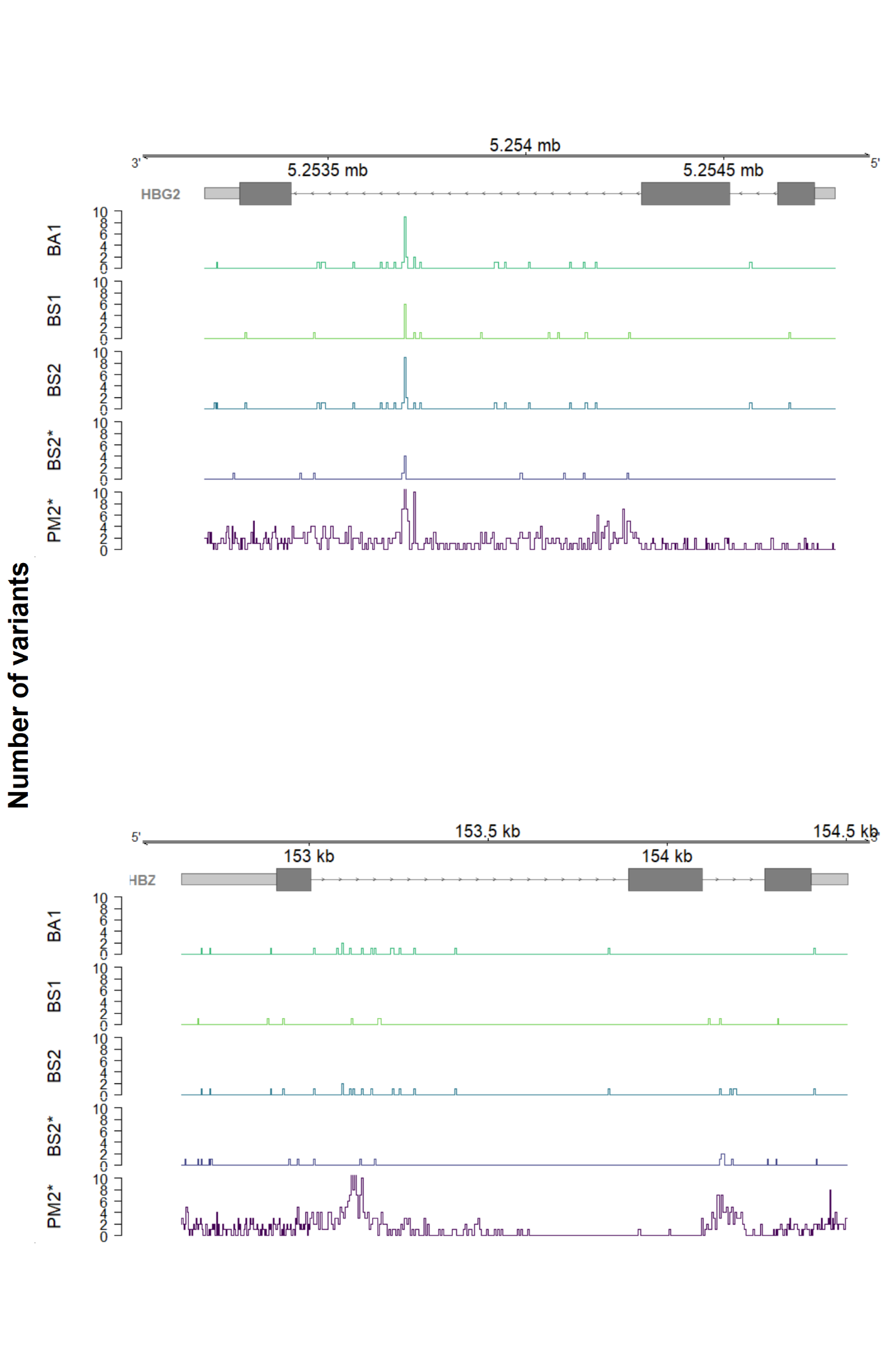

**Supplementary Figure 3. Genomic distribution of globin gene variants meeting ACMG/AMP criteria in gnomAD v4.1.** Density tracks show quality-passing variants from the gnomAD v4.1 joint dataset across *HBA2*, *HBA1*, *HBB*, *HBD*, *HBE1*, *HBG1*, *HBG2*, and *HBZ*, stratified by ACMG/AMP criteria met (**BA1**, green; **BS1**, light green; **BS2**, blue; **BS2***, lavender; **PM2***, purple, where * denotes the _Supporting strength level, corresponding to BS2_Supporting and PM2_Supporting). Gene models and genomic coordinates are shown above each panel; where horizontal lines denote introns, light grey boxes denote untranslated regions, and dark grey boxes denote coding sequences. Gene tracks are displayed using 3-bp exonic bins and 5-bp intronic bins.
