## Supplementary Table for "Analyzing globin gene variation in gnomAD: implications for variant interpretation in hemoglobinopathies"

**Supplementary Table 1. Variants from the gnomAD v4.1 release that passed quality assessment and met conflicting ACMG/AMP criteria.**

| **Gene** | **Position** | **Reference allele** | **Alternative allele** | **ClinVar annotation** | **Population maximum filtering allele frequency** | **Total allele frequency** | **BS1** | **BS2** | **PM2** |
| --- | --- | --- | --- | --- | --- | --- | --- | --- | --- |
| *HBB* | Chr11:5225469-5225469 | A | T | Uncertain significance | 0.00142012 | 0.00008863370 | Met | Not met | Met |
| *HBB* | Chr11:5225763-5225763 | C | A | Benign/Likely benign | 0.00166359 | 0.00009864220 | Met | Not met | Met |
| *HBB* | Chr11:5226646-5226646 | G | T | Uncertain significance | 0.00156877 | 0.00005761530 | Met | Not met | Met |
| *HBD* | Chr11:5233045-5233045 | C | T | Benign/Likely benign | 0.00118245 | 0.00008302200 | Met | Not met | Met |
| *HBE1* | Chr11:5269517-5269517 | G | A | Not available | 0.00104667 | 0.00004708800 | Met | Not met | Met |
| *HBE1* | Chr11:5269791-5269791 | A | G | Not available | 0.00124429 | 0.00008331440 | Met | Not met | Met |
| *HBG1* | Chr11:5248923-5248923 | T | C | Not available | 0.00121129 | 0.00007374730 | Met | Not met | Met |
| *HBG2* | Chr11:5253698-5253698 | G | GTGCACACACACACACACA | Not available | 0.00135199 | 0.00009661550 | Met | Not met | Met |
| *HBG2* | Chr11:5253889-5253889 | C | T | Not available | 0.00133795 | 0.00007885810 | Met | Not met | Met |
| *HBB* | Chr11:5226434-5226434 | C | T | Benign/Likely benign | 0.00055887 | 0.00009607180 | Not met | Met | Met |
| *HBB* | Chr11:5227003-5227005 | CAG | C | Pathogenic/Likely pathogenic | 0.00022786 | 0.00002358730 | Not met | Met | Met |
| *HBG1* | Chr11:5248376-5248376 | A | C | Not available | 0.00029254 | 0.00002725850 | Not met | Met | Met |
| *HBG1* | Chr11:5249015-5249015 | T | C | Not available | 0.00000000 | 0.00007021880 | Not met | Met | Met |
| *HBG1* | Chr11:5249173-5249173 | T | A | Not available | 0.00005548 | 0.00003744760 | Not met | Met | Met |
| *HBG1* | Chr11:5249235-5249235 | C | T | Not available | 0.00022732 | 0.00002809480 | Not met | Met | Met |
| *HBG1* | Chr11:5249253-5249253 | C | T | Not available | 0.00004540 | 0.00006258170 | Not met | Met | Met |
| *HBG1* | Chr11:5249259-5249259 | C | T | Not available | 0.00000573 | 0.00000844960 | Not met | Met | Met |
| *HBG1* | Chr11:5249288-5249288 | G | A | Not available | 0.00006229 | 0.00000986295 | Not met | Met | Met |
| *HBG1* | Chr11:5249326-5249326 | C | T | Not available | 0.00000849 | 0.00000904443 | Not met | Met | Met |
| *HBG1* | Chr11:5249327-5249327 | G | A | Not available | 0.00002030 | 0.00002580500 | Not met | Met | Met |
| *HBG1* | Chr11:5249358-5249358 | C | T | Not available | 0.00000393 | 0.00000805065 | Not met | Met | Met |
| *HBG1* | Chr11:5249388-5249388 | C | T | Not available | 0.00000464 | 0.00000705577 | Not met | Met | Met |
| *HBA1* | Chr16:177078-177078 | C | G | Uncertain significance | 0.00009792 | 0.00000910486 | Not met | Met | Met |
| *HBA1* | Chr16:177340-177340 | C | T | Pathogenic/Likely pathogenic | 0.00007740 | 0.00003099470 | Not met | Met | Met |
| *HBA1* | Chr16:177482-177482 | C | G | Not available | 0.00006051 | 0.00001795870 | Not met | Met | Met |
| *HBA1* | Chr16:177483-177485 | GTA | G | Not available | 0.00006081 | 0.00001868950 | Not met | Met | Met |
| *HBA1* | Chr16:177488-177488 | C | CTT | Not available | 0.00010144 | 0.00002384860 | Not met | Met | Met |
| *HBA1* | Chr16:177490-177491 | CG | C | Not available | 0.00011581 | 0.00002390380 | Not met | Met | Met |
| *HBA2* | Chr16:173363-173363 | G | A | Uncertain significance | 0.00031376 | 0.00003060530 | Not met | Met | Met |
| *HBA2* | Chr16:173385-173385 | C | T | Not available | 0.00020320 | 0.00001683130 | Not met | Met | Met |
| *HBA2* | Chr16:173395-173395 | G | T | Not available | 0.00006728 | 0.00000560846 | Not met | Met | Met |
| *HBA2* | Chr16:173448-173448 | G | C | Not available | 0.00039848 | 0.00008501470 | Not met | Met | Met |
| *HBA2* | Chr16:173497-173497 | C | A | Uncertain significance | 0.00069275 | 0.00004731210 | Not met | Met | Met |
| *HBA2* | Chr16:173546-173546 | C | G | Benign/Likely benign | 0.00037496 | 0.00007713350 | Not met | Met | Met |
| *HBA2* | Chr16:173576-173576 | C | T | Not available | 0.00006555 | 0.00000559647 | Not met | Met | Met |
| *HBA2* | Chr16:173580-173580 | C | A | Uncertain significance | 0.00022124 | 0.00001492360 | Not met | Met | Met |
| *HBA2* | Chr16:173585-173585 | C | T | Benign/Likely benign | 0.00002980 | 0.00002051860 | Not met | Met | Met |
| *HBA2* | Chr16:173608-173608 | C | T | Benign/Likely benign | 0.00004579 | 0.00000559600 | Not met | Met | Met |
| *HBA2* | Chr16:173618-173618 | C | G | Not available | 0.00010223 | 0.00002424640 | Not met | Met | Met |
| *HBA2* | Chr16:173692-173692 | A | G | Pathogenic/Likely pathogenic | 0.00021222 | 0.00003825760 | Not met | Met | Met |
| *HBZ* | Chr16:153124-153124 | T | TGGGGAGGGGACAGTGGGGAGAGGACAGTAAGGAGGGGACCATGGGGAGGACACAGG | Not available | 0.00003669 | 0.00006094890 | Not met | Met | Met |
| Data sources: gnomAD (accessed 28 May 2025) and ClinVar (accessed 10 October 2025). | | | | | | | | | |
